# A deletion in *GDF7* is associated with a heritable forebrain commissural malformation concurrent with ventriculomegaly and interhemispheric cysts in cats

**DOI:** 10.1101/2020.05.12.091686

**Authors:** Yoshihiko Yu, Erica K. Creighton, Reuben M. Buckley, Leslie A. Lyons, 99 Lives Consortium

**Affiliations:** Department of Veterinary Medicine and Surgery, College of Veterinary Medicine, University of Missouri, Columbia, MO, 65211, USA; Laboratory of Veterinary Radiology, Nippon Veterinary and Life Science University, Musashino, Tokyo, 180-8602, Japan

**Keywords:** feline, *Felis catus*, brain malformation, BMP12, neurodevelopment, genetics, genomics, Mendelian traits, genome-wide association study, whole genome sequencing

## Abstract

An inherited neurologic syndrome in a family of mixed-breed Oriental cats has been characterized as forebrain commissural malformation concurrent with ventriculomegaly and interhemispheric cysts. However, the genetic basis for this autosomal recessive syndrome in cats is unknown. Forty-three cats were genotyped on the Illumina Infinium Feline 63K iSelect DNA Array and used for analyses. Genome-wide association studies, including a sib-transmission disequilibrium test, a case-control association analysis, and homozygosity mapping, identified a critical region on cat chromosome A3. Short-read whole genome sequencing was completed for a cat trio segregating with the syndrome. A homozygous 7 bp deletion in *growth differentiation factor 7* (*GDF7*) (c.221_227delGCCGCGC [p.Arg74Profs]) was identified in affected cats by comparison to the 99 Lives Cat variant dataset, validated using Sanger sequencing, and genotyped by fragment analyses. This variant was not identified in 192 unaffected cats in the 99 Lives dataset. The variant segregated concordantly in an extended pedigree. Obligate carrier cats were heterozygous. In mice, *GDF7* mRNA is expressed within the roof plate when commissural axons initiate ventrally-directed growth. This finding emphasizes the importance of *GDF7* in the neurodevelopmental process in the mammalian brain. A genetic test can be developed for use by cat breeders to eradicate this variant.

## 1. Introduction

Congenital brain malformations in humans are caused by genetic variants, *in utero* infection, or other environmental factors. Dogs and cats are also occasionally diagnosed with congenital brain malformations (Reviewed in [1]), which are noted as breed predispositions, familial aggregations, and sporadic cases [2-6]. Congenital hydrocephalus is common in toy and brachycephalic dog breeds, such as the Maltese, Yorkshire terrier, Chihuahua, toy poodle, and pug dogs [7]. Widespread in Cavalier King Charles Spaniels, Chiari-like malformation is a common cause of foramen magnum obstruction and results in the secondary syringomyelia in dogs, characterized by the mismatch of size between the brain and the skull [8].

Similarly, high grades of brachycephaly in cats are also associated with malformations of the calvarial and facial bones, as well as dental malformations or respiratory abnormalities [9-12]. Burmese cats have a familial craniofacial malformation with meningoencephalocele [13] that is caused by *ALX Homeobox 1* (*ALX1)* variant [14]. However, feline brain malformations with (suspected) idiopathic nature are mostly reported as sporadic events [15-19]. Overall, the genetic factors contributing to brain (mal)formation and structural congenital brain disease in dogs and cats are largely unknown.

An extended family of mixed-breed cats derived from the Oriental breed has been characterized clinically and histopathologically with forebrain commissural malformation concurrent with ventriculomegaly and interhemispheric cysts [20]. These cats have small rounded ear pinnae and doming of the head (**Figure 1**). The forebrain malformations include dysgenesis of the septum pellucidum, interthalamic adhesion, and all the midline commissures, excluding the rostral white commissure, as well as hippocampal hypoplasia. Clinical symptoms include mild generalized ataxia when walking and mild to marked postural reaction deficits, although cranial nerve examination and segmental reflexes are within normal limits. All the cats with neurological signs have midline and limbic structure abnormalities, dilated ventricles, and hemispheral cysts with or without a suprapineal cyst. These findings resemble a mild variant of holoprosencephaly (HPE) in human (OMIM: 236100 and others). Although variations in the severity of the forebrain commissural malformation were seen, most affected cats are hydrocephalic. This malformation was identified in a breeding program designed to propagate a small, rounded ear. No chromosomal abnormalities are noted in a karyotypic analysis of the cats. Segregation analysis suggests an autosomal recessive mode of inheritance; however, the causal variant remains unknown [20].

**Figure 1.**
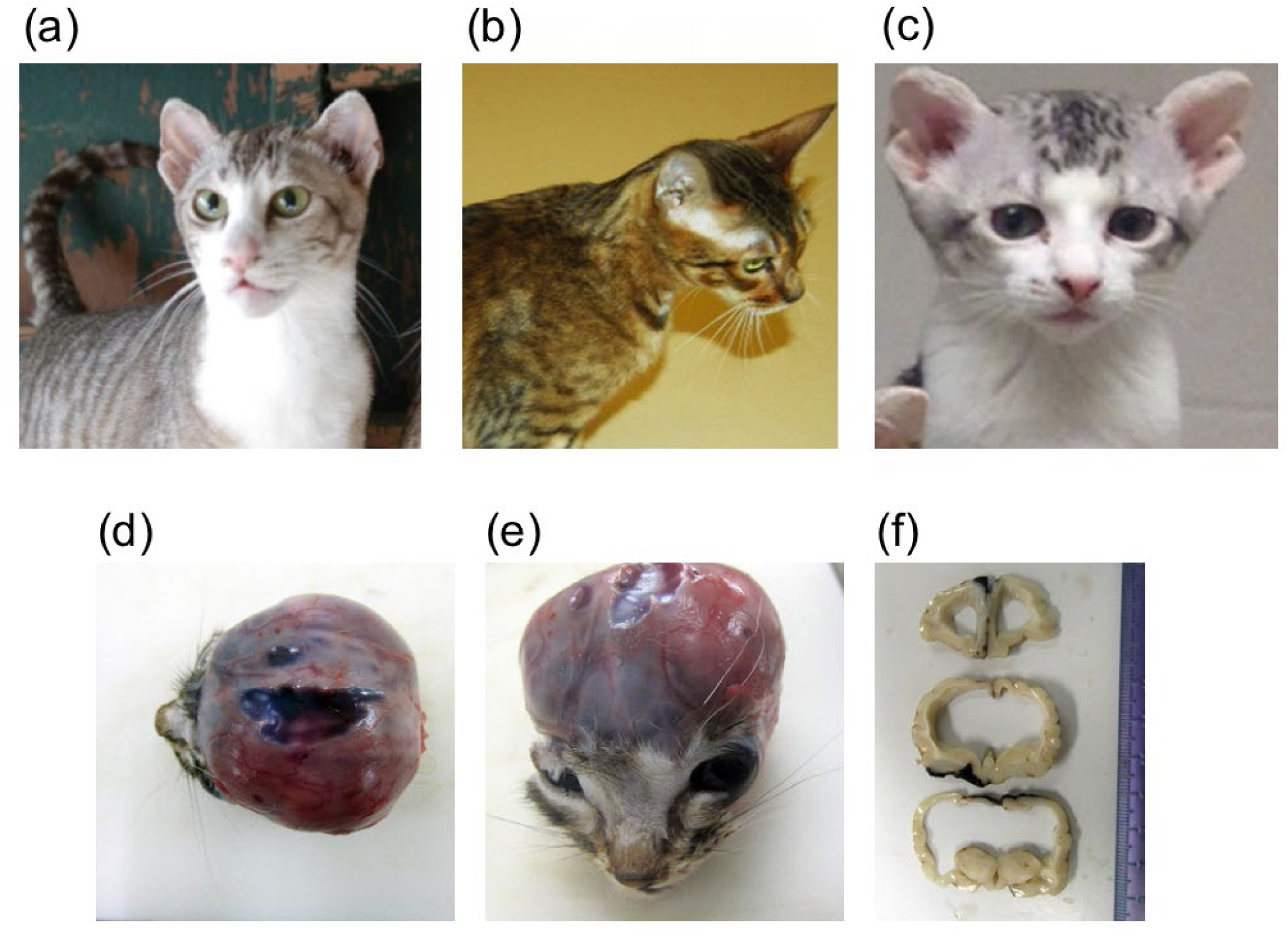
Domestic cats with feline forebrain commissural malformation in cats. Note the abnormal presentation of the pinnae used to determine affection status. (a) Frank – affected sire (left). (b) Camilla – carrier queen. (c) Bobble – affected offspring. These three cats (a–c) were whole genome sequenced. (d) Transverse plane of T2-weighted magnetic resonance imaging of an affected cat at the level of the thalamus. Severe ventriculomegaly, thinning of the cerebral parenchyma, and midline structure deficits are seen. A part of the parietal lobe is deficient. (e) Mid-sagittal plane of T2-weighted magnetic resonance imaging of an affected cat (the same cat as [d]). Midline structure deficits are recognized. Note that the spinal cord is formed normally. Interhemispheric cysts are also seen at the rostrotentorial region and the quadrigeminal cistern. Due to the presence of cysts, cerebellar herniation is seen. (f) Gross dorsal view of the dissected head at necropsy. The skin was removed, and the skull was exposed. (g) Transverse sections of formalin-fixed brain tissue at the level of frontal lobe and thalamus. Severe ventriculomegaly, thinning of the cerebral parenchyma, and midline structure deficits are seen. Note that a cat whose magnetic resonance imaging of (d) and (e) are presented here is different from cats whose gross pathological pictures of (f) and (g) are provided here.

Genome-wide association studies (GWAS), using a sib-transmission disequilibrium test (sib-TDT) and case-control analysis, and homozygosity mapping were conducted to detect an associated genomic region for the syndrome using genotypes from a feline single nucleotide polymorphism (SNP) DNA array [21]. Whole genome sequencing (WGS) was conducted for a cat trio segregating for the syndrome to refine the location and identify candidate variants.

## 2. Materials and Methods

### 2.1. Sampling and pedigree

All procedures were performed with an approved University of Missouri (MU) Institutional Animal Care and Use Committee protocol (ACUC protocol # 8292). Four affected and two carrier cats were donated and housed at the MU colony for controlled breeding. Additional buccal swab and cadaver samples from an external breeding program were provided voluntarily by the breeder/owner (N = 129). DNA samples were extracted using DNeasy Blood & Tissue Kit (Qiagen, Valencia, CA, USA). The quality of the DNA samples was visualized and confirmed by agarose gel electrophoresis. DNA samples whose concentration was insufficient were whole genome amplified using the REPLI-g Mini Kit (Qiagen). The relationship of the ascertained cats was confirmed using short tandem repeat (STR) markers as previously described [22]. Parentage analysis was performed using the computer program Colony [23,24] (data not shown). Clinical and histopathological features of the syndrome were characterized previously [20]. Although some cats were phenotyped based on magnetic resonance imaging and/or histopathology, most cats were assumed to have the brain malformation based on the ear morphology since clinically healthy cats had elongate (normal) ears and clinically affected cats had the small, rounded ear type [20] (**Figure 1**). Images or cadavers of cats were not always available.

### 2.2. DNA array genotyping

Fifty-two genomic DNA samples (∼600 ng each) were submitted to GeneSeek (Neogene, Lincoln, NE, USA) for SNP genotyping on the Illumina Infinium Feline 63K iSelect DNA Array (Illumina, San Diego, CA, USA) [21]. Since SNP positions were based on an early assembly of the cat genome [25], the SNPs were relocalized to the latest feline genome assembly, Felis_catus_9.0 [26]. Quality control of the SNP data was performed using PLINK (v1.07) [27]. The following criteria were applied: (i) individuals with genotyping success rate of <80% were removed (--mind 0.2), (ii) SNP markers with a genotyping rate <80% were removed (--geno 0.2), and, (iii) SNPs with a minor allele frequency of 0.05 or less were removed (--maf 0.05). Furthermore, SNPs that were previously reported to have missing ≥ 10% of genotypes and Mendelian errors [21], and that remained after quality control were excluded.

### 2.3. Genome-wide association studies

After the SNP pruning described above, GWAS were conducted using PLINK. Sibling transmission disequilibrium test (sib-TDT) [28] was performed using the DFAM procedure in PLINK (--dfam). This method implements sib-TDT and also includes unrelated individuals in the analysis. A case-control association analysis was performed (--assoc). The genomic inflation factor was calculated using the function (--adjust). Multi-dimensional scaling (MDS) analysis was conducted (--genome) and MDS plots were generated to visualize the population stratification, using PLINK and R software (version 3.3.3; R Foundation for Statistical Computing, Vienna, Austria), respectively. A quantile-quantile (QQ) plot was created using R. Genome-wide significance for both analyses was determined using 100,000 permutations (--mperm 100000). Manhattan plots from the sib-TDT, case-control association, and permutation analyses were generated using R. The MDS plot was used to reselect cats to minimize stratification between cases and controls for secondary case-control association analysis.

### 2.4. Haplotype analysis

An approximately 6 Mb region surrounding highly associated SNPs was extracted including 81 SNPs, from SNP chrA3.163737349 at chromosome position A3: 123,014,546 to SNP chrA3.156620632 at chromosome position A3: 128,837,125. The haplotype boundaries were visually confirmed using Haploview (version 4.2) [29].

### 2.5. Homozygosity analysis

Homozygosity analysis was performed using PLINK. SNPs within a 1000 kb window, containing at least 25, were investigated for runs of homozygosity (--homozyg-window-kb 1000, -- homozyg-snp 25). In each window, five missing genotypes (20%) and a single heterozygote (2%) were tolerated (--homozyg-window-missing 5, --homozyg-window-het 1). The threshold of homozygosity match was set as 0.99 (--homozyg- match 0.99). A homozygous block was characterized by five SNPs (∼200 - 250 kb). Consensus homozygosity blocks were identified as overlaps between individual homozygosity blocks (--consensus-match, --homozyg-group).

### 2.6. Whole genome sequencing

A trio of cats including an affected sire, a carrier dam, and an affected offspring was selected for WGS as part of the 99 Lives Cat Genome Sequencing Initiative (http://felinegenetics.missouri.edu/99lives). These cats were produced at the MU colony; thus, parentage was known. DNA extraction and library preparation were conducted as previously described [30]. A minimum of 4 µg genomic DNA was submitted for WGS to the MU DNA Core Facility. Two PCR-free libraries with insertion sizes of 350 bp and 550 bp were constructed for each cat using the TruSeq DNA PCR Free library preparation kit (Illumina). The Illumina HiSeq 2000 (Illumina) was used to generate sequence data.

Sequence reads were mapped to the latest feline genome assembly, Felis_catus_9.0, and processed as previously described [26]. Briefly, read mapping was conducted with Burrows-Wheeler Aligner (BWA) version 0.7.17 [31]. Duplicates were marked using Picard tool MarkDuplicates (http://broadinstitute.github.io/picard/). Potential insertions or deletions (indels) realignment was performed using the Genome Analysis Tool Kit (GATK version 3.8) [32] IndelRealigner. Variants were called using GATK HaplotypeCaller in gVCF mode [33]. VarSeq v2.0.2 (Golden Helix, Bozeman, MT) was used to annotate variants with Ensembl 99 gene annotations and identify variants unique to the trio cats and absent from 192 unaffected unrelated domestic cats. Exonic variants were extracted from the dataset, including variants 21 bp flanking the exons. Candidate variants segregating across the trio were visualized using Integrative Genomics Viewer (IGV) [34].

### 2.7. Variant validation and genotyping

PCR and Sanger sequencing were performed to validate the 7 bp deletion in the candidate gene *GDF7* for cats that were submitted to WGS. The primer sequences were; Forward primer: 5’- AGCGACATCATGAACTGGTG -3’, Reverse primer: 5’- CCACGGAGCCCATGGACC -3’. PCR was performed using AccuPrime GC-Rich DNA Polymerase (Invitrogen, Carlsbad, CA, USA). PCR was performed following the manufacturer’s instructions with the annealing temperature of 61°C and 35 cycles. PCR amplicon was purified using QIAquick Gel Extraction Kit (Qiagen) or using ExoSAP-IT PCR Product Cleanup Reagent (Thermo Fisher Scientific, Waltham, MA, USA). Sanger sequencing was conducted at the MU DNA Core Facility using an Applied Biosystems 3730xl DNA Analyzer (Applied Biosystems, Foster City, CA, USA) with BigDye Terminator v3.1 Cycle Sequencing Kit (Applied Biosystems).

Fragment analysis was conducted for population screening. PCR conditions and reagents were as above, except the forward primer was fluorescein amidite [FAM] labeled at the 5’ end. Fragment analysis was conducted at the MU DNA Core Facility using an Applied Biosystems 3730xl DNA Analyzer (Applied Biosystems). The expected wildtype fragment size was 294 bp, while the mutant fragment size was expected as 287 bp. Amplicons were analyzed using STRand [35].

## 3. Results

### 3.1. Pedigree and genotyping

Using 18 STRs, parentage for 69 of 129 cats was determined with a high likelihood using the COLONY software [23,24] (data not shown), producing a pedigree of 79 cats (**Figure S1**). For GWAS, 52 cats were selected using owner provided and pedigree information, including 26 cases, and 26 controls in which 43 cats were included in the pedigree (**Figure S1**). Cat DNA samples were genotyped on Feline 63K SNP array (**File S1**). Selection criteria for genotyping focused on cats that were as unrelated as possible. Nine cats with call rates below 80% were removed, and 478 SNPs were removed with missingness rates > 20%. An additional 22,297 SNPs were also removed with minor allele frequencies < 0.05. After filtering, 20 cases and 23 controls remained with a genotyping rate of 0.977 across 40,263 SNPs. Furthermore, 377 SNPs were excluded due to missing ≥ 10% of genotypes and Mendelian errors previously reported [21]. The GWAS was conducted with 39,891 SNPs.

### 3.2. Association Studies

Sib-TDT was conducted on the pedigree formed by the 20 cases and 23 controls. After permutation testing, no SNPs were significant; however, 9 SNPs with the highest, the second-highest, or the third-highest association were localized to cat chromosome A3:123,055,238 – 128,667,138 on the Felis_catus_9.0, extending approximately 5.6 Mb (**Table 1**). The result of the sib-TDT analysis was presented as a Manhattan plot (**Figure 2a**). In the initial case-control association analysis, 65 SNPs had genome-wide significance and were located cat chromosome A3: 116,714,934– 129,668,450, extending ∼13.0 Mb and C1: 105,429,018 – 115,412,315, extending ∼10.0 Mb (**Table 1**). However, the genomic inflation factor was 1.89; thus, the MDS plot (**Figure S2**) was used to reselect cases and controls for the analysis. A second case-control association analysis was performed with 14 cases and 9 controls, and the genomic inflation factor was reduced to 1. Seventeen SNPs showed genome-wide significance and were located cat chromosome A3: 119,105,247 – 129,372,537, encompassing ∼10.3 Mb (**Figure 2b, Table 1**). This chromosome A3 region encompassed the entire region suggested by the sib-TDT and was within the initial case-control association analysis.

**Table 1.** SNPs associations for hydrocephalus cats in the sib-TDT and case-control association analyses.

| SNP | chr | Position | sib-TDT | Case-control (Initial) | Case-control (2nd) |
| --- | --- | --- | --- | --- | --- |
| chrA3.164724433 | A3 | 123353491 | >0.6 | 0.00011 | 0.00449 |
| chrA3.164567500 | A3 | 122318611 | >0.6 | 0.00006 | 0.00449 |
| chrA3.164340161 | A3 | 122513677 | >0.6 | 0.00011 | 0.00449 |
| chrA3.164113252 | A3 | 122698750 | >0.6 | 0.00006 | 0.00449 |
| chrA3.163320257 | A3 | 123353491 | 0.5639 | 0.00006 | 0.00449 |
| chrA3.162970354 | A3 | 123644765 | >0.6 | 0.00006 | 0.00449 |
| chrA3.162343840 | A3 | 124176474 | >0.6 | 0.00036 | 0.00449 |
| chrA3.158624618 | A3 | 127189752 | >0.6 | 0.00011 | 0.00449 |
| chrA3.159621145 | A3 | 126377299 | 0.5639 | 0.00005 | 0.0082 |
| chrA3.162413594 | A3 | 124100380 | >0.6 | 0.00136 | 0.01369 |
| chrA3.156826206 | A3 | 128667138 | 0.5639 | 0.00006 | 0.01685 |
| chrA3.156620632 | A3 | 128837125 | >0.6 | 0.00011 | 0.01685 |
| chrA3.155936886 | A3 | 129372537 | >0.6 | 0.00037 | 0.01685 |
| chrA3.168960567 | A3 | 119105247 | >0.6 | 0.01283 | 0.02607 |
| chrA3.168031908 | A3 | 119810207 | >0.6 | 0.02638 | 0.02607 |
| chrA3.167492986 | A3 | 120088757 | >0.6 | 0.00605 | 0.02607 |
| chrA3.167322483 | A3 | 120215597 | >0.6 | 0.02922 | 0.02607 |
| chrA3.162621987 | A3 | 123934341 | 0.5521 | 0.00136 | >0.05 |
| chrA3.163679766 | A3 | 123055238 | 0.5639 | 0.00035 | >0.05 |
| chrA3.161984351 | A3 | 124475589 | 0.5639 | 0.0002 | >0.05 |
| chrA3.161943004 | A3 | 124509146 | 0.5639 | 0.0002 | >0.05 |
| chrA3.161399869 | A3 | 124945294 | 0.5639 | 0.00197 | >0.05 |
| chrA3.160673309 | A3 | 125511595 | 0.5639 | 0.00021 | >0.05 |

\*SNP IDs are based on an early cat genome assembly [23] †Positions based on current cat genome assembly [24].

**Figure 2.**
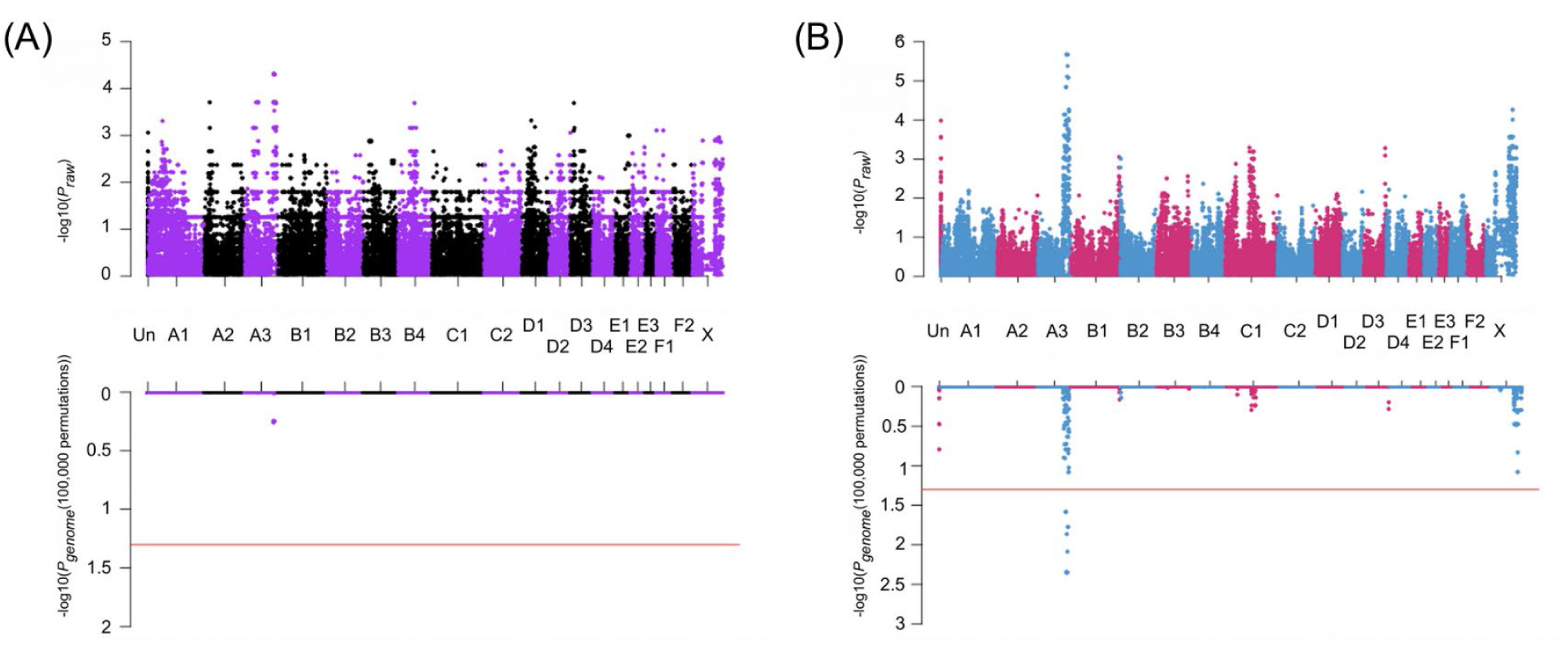
Manhattan plot of the GWAS analyses for feline forebrain commissural malformation in cats. Cats (20 cases and 23 controls) were genotyped on the Illumina Infinium Feline 63K iSelect DNA Array (Illumina, San Diego, CA, USA). In both panels of (a) and (b), upper plots exhibit the *P*_*raw*_ value of the analysis, while the lower exhibits the *P*_*genome*_ values after 100,000 permutations. Red horizontal lines indicate genome-wide significance (*P*_*genome*_ = 0.05, -log_10_ = 1.3). (a) Sib-TDT analysis. Genome-wide significance was not achieved (b) Case-control association analysis. Significant association is localized to chromosome A3 for 17 SNPs. Genomic inflation was 1.

### 3.3. Haplotype analysis

The 6 Mb region covering chrA3: approximately 123 – 129 Mb and encompassing the overlapped region with identified in GWAS, was visually inspected for common haplotypes, using Haploview. In affected cats, a single unique haplotype encompassing 4,265 kb was identified with 95% frequency using the solid spine of LD method in Haploview. Considering two cats had 82.7% and 91.4% genotyping rate, one cat had 98.8%, and the others had 100% genotyping rate in this area, a few missing produced the remaining haplotypes (**Figure S3, File S2**).

### 3.4. Homozygosity analysis

Homozygosity mapping was performed on 20 cases and 23 controls. The homozygosity analysis identified the same location on chromosome A3 in 18 of 20 affected cats, excluding the same cases with A3: 125,601,560 – 127,684,693, spanning approximately 2.1 Mb and no unaffected cats were homozygosity (**Table S1**). The region was identified by the two genome-wide association analyses (**Table 1**). Although other ROHs were identified, none were specific to cases or as extensive.

### 3.5. Whole genome sequencing

Cat genomes have been submitted to the NCBI short read archive under BioProject: PRJNA528515; Accessions PRJNA343385; SRX2654400 (Sire), SRX2654398 (dam) and SRX2654399 (offspring). Genome sequence analyses and variant calling for the 99 Lives project has been previously described [36]. Approximately 2.5 million variants were ascertained across 195 cats in the exonic portion of the dataset, which included 21 bp of exon flank sequence. No candidate genes were identified on cat chromosome A3 during the initial analysis when considering the sire and offspring to be homozygous affected and the queen as an obligate carrier for an alternative allele (**Table 2**). Only an intergenic variant (C1:106,990,675) and an intronic variant in *sperm antigen with calponin homology and coiled-coil domains* (*SPECC1)* (E1:9,973,078) met the segregation criteria. Using relaxed constraints, where affected cats were allowed to also be considered as carriers, four more variants were identified (C1:96,095,693, C1:96,839,645 and D2:33,368,378) with only one variant located within the critical region and also in a gene coding region (**Table 2**). This variant was a 7 bp deletion in the coding region of *GDF7* (c.221_227delGCCGCGC [p.Arg74Profs*17]) at position A3:127002233 (ENSFCAT00000063603). The variant was identified as homozygous in the affected sire, heterozygous in the obligate carrier dam, heterozygous in the affected offspring and absent from the other 192 domestic cats. Although each cat in the trio had an average of ∼30× genome coverage, the sire had 18× coverage within the region, the queen had ∼14× coverage with 7 reads per allele, and the affected offspring had ∼16× coverage with only one of the reads representing the reference allele, likely misrepresenting the offspring as heterozygous (**Figure 3**). The affected cat was confirmed as a homozygote for the alternate allele by genotyping. The *GDF7* variant was predicted to cause a truncated protein with 89 amino acids, while the wildtype protein has 455 amino acids (**Figure S4**). Feline GDF7 amino acid sequence is predicted to be 86.2%, 90.1%, 84.6%, 77.8%, and 77.2% identical to human, horse, cow, rat, and mouse, respectively (**Figure S4**). In addition, comparison of the *GDF7* locus between the Felis_catus_9.0 and Felis_catus_8.0 genome assemblies, revealed the region containing the *GDF7* candidate variant is absent from Felis_catus_8.0 assembly, indicating the importance of the updated reference genome for trait discovery.

**Table 2.**
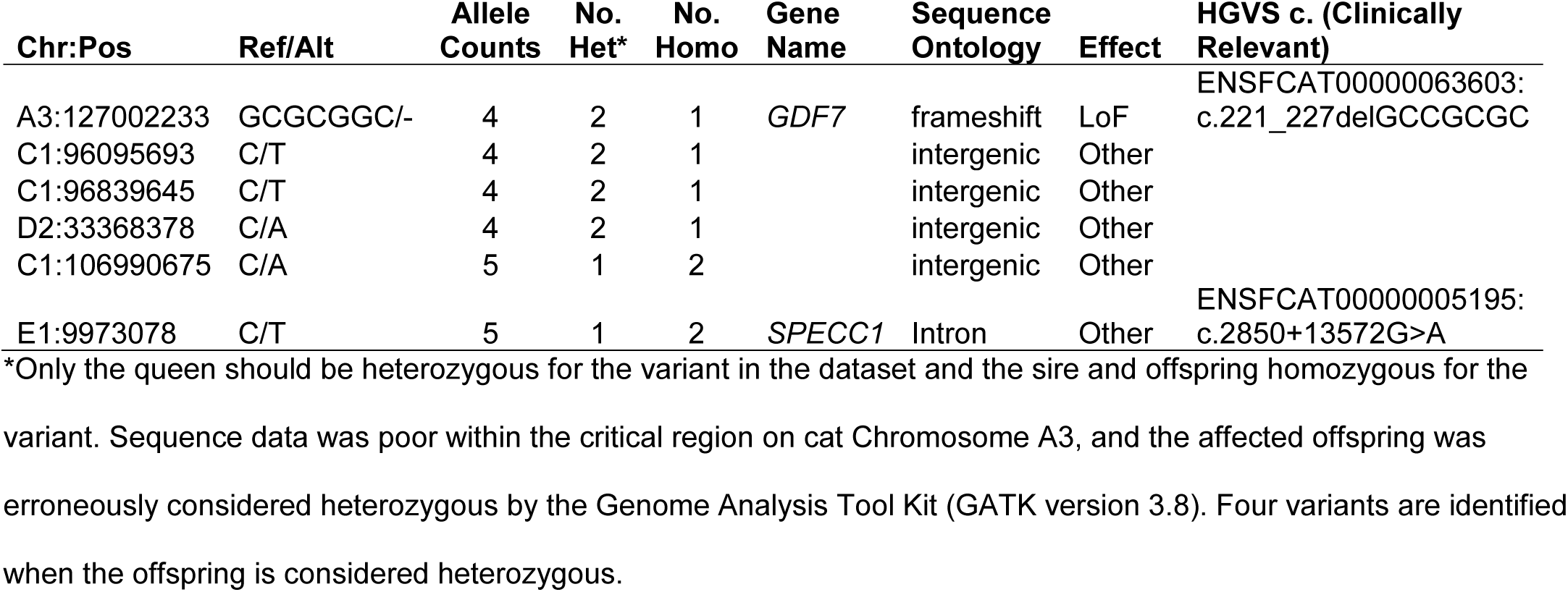
Variants identified in 99 Lives whole genome sequence dataset considering segregation within the trio.

| Chr:Pos | Ref/Alt | Allele Counts | No. Het* | No. Homo | Gene Name | Sequence Ontology | Effect | HGVS c. (Clinically Relevant) |
| --- | --- | --- | --- | --- | --- | --- | --- | --- |
| A3:127002233 | GCGCGGC/- | 4 | 2 | 1 | *GDF7* | frameshift | LoF | ENSFCAT00000063603:<br>c.221_227delGCCGCGC |
| C1:96095693 | C/T | 4 | 2 | 1 |  | intergenic | Other |  |
| C1:96839645 | C/T | 4 | 2 | 1 |  | intergenic | Other |  |
| D2:33368378 | C/A | 4 | 2 | 1 |  | intergenic | Other |  |
| C1:106990675 | C/A | 5 | 1 | 2 |  | intergenic | Other |  |
| E1:9973078 | C/T | 5 | 1 | 2 | *SPECC1* | Intron | Other | ENSFCAT00000005195:<br>c.2850+13572G>A |

\*Only the queen should be heterozygous for the variant in the dataset and the sire and offspring homozygous for the
variant. Sequence data was poor within the critical region on cat Chromosome A3, and the affected offspring was
erroneously considered heterozygous by the Genome Analysis Tool Kit (GATK version 3.8). Four variants are identified
when the offspring is considered heterozygous.

**Figure 3.**
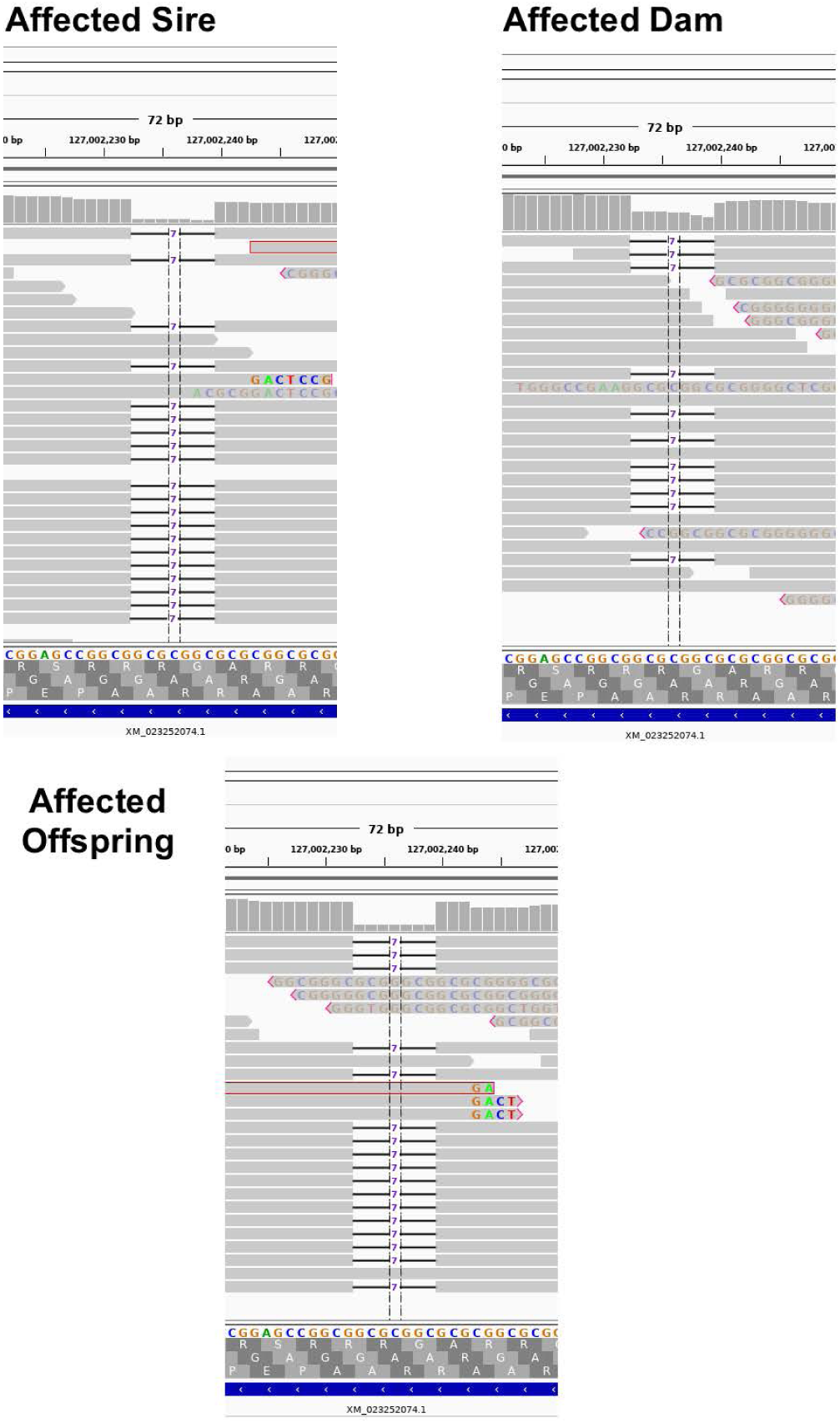
Depiction of the whole genome sequence reads using the Integrated Genome Viewer (IGV) for the *GDF7* variant. Grey horizontal bars represent individual reads, while grey vertical bars at the top of each sub figure represent depth of coverage. Notice in affected individuals, coverage is close to zero in the deleted region, while in the carrier sequencing coverage is approximately 50%. Also, in affected individuals, reads with high numbers of mismatches are indicative of misidentification of an indel and tend to occur near the ends of reads.

### 3.6. Variant validation and genotyping

Sanger sequencing was performed to confirm the identified *GDF7* c.221_227delGCCGCGC in affected and obligate carrier cats, including the cats in the WGS trio. The 7 bp deletion in *GDF7* was screened in 25 affected, 39 unaffected, and two cats with unknown phenotype in the extended pedigree using fragment analysis (**Figure 4, Table S1)**. Both unknown cats were homozygous for the variant allele. Overall, 13 of 14 suspected wildtype cats in the extended pedigree were concordant, and one cat genotyped as a heterozygote. Of 25 suspected carriers, 23 genotyped as heterozygote and two as wildtype normal. Of 22 suspected affected cats, 20 genotyped as homozygous for the variant, one as heterozygous and one as wildtype normal.

## 4. Discussion

Brain malformations are occasionally identified in veterinary practice. However, little is known about the genetic causes and interactions for brain malformation. Due to the health concerns associated with breed development, particularly in dog breeds [37,38], many breeders have become more vigilant to health-associated consequences of selection based on morphological phenotypes. In an effort to develop a breed of cats having similar phenotypes to a tiger, including a small rounded ear, a mixed breed cat derived from the Oriental cat breed was discovered to have small rounded ears, and hence, was used as a foundation sire for a breeding program. Outcross and backcross breeding indicated the phenotype was autosomal recessive [20]. However, an MRI examination of a kitten with the desired ear phenotype, which had an accidental head injury from a fall, indicated the presence of congenital hydrocephalus. Additional MRIs of the breeding stock suggested all cats with the ear phenotype had congenital hydrocephalus without the findings of increased intracranial pressure. As a result of the potentially harmful impacts associated with the trait, the breeder promptly discontinued the breeding program and altered subsequent cats. However, some carriers for the trait had already been adopted for other breeding programs. A group of affected cats were presented to the researchers for pathological and genetic studies. Various brain midline deficits concurrent with ventriculomegaly and interhemispheric cysts are present in the cats with the small rounded pinnae [20]. Sample collection from the cats in the owner’s breeding program and cats from controlled breeding within the university colony supported the genetic investigation of the abnormal brain development and mode of inheritance. GWAS, homozygosity mapping, and WGS were conducted to detect the underlying causal variant for the brain malformation in the cats.

Most of the cat samples had been archived as frozen cadavers by the breeder and later provided to the researchers. Because of poor documentation as to relationships and disease status, a pedigree was established by determining parentage using STRs, ages, and gender of the cats and from interviews with the breeder. Ear phenotypes, which were used as a proxy for disease, were difficult to determine from frozen cadavers. Due to the significant inbreeding and backcrossing required to maintain the phenotype, 18 STRs were often insufficient to determine parentage. However, some known breedings were available from the university colony. Overall, an extended pedigree was developed and was expected to be sufficient for GWAS and WGS investigations for the causal variant.

GWAS, including sib-TDT and case-control association analyses, were performed to localize the syndrome to the cat genome. A sib-TDT was conducted as maternal DNA samples for most trios were unavailable; therefore, a TDT for family trio design was likely to lack sufficient power. An association on cat chromosome A3, where the highest associated SNPs spanned the region of ∼123.1 to 128.7 Mb, was identified but without genome-wide significance after permutation. Additionally, a case-control association analysis was also performed with the same cats, and an association was identified in the same region of chromosome A3. Genome-wide significance was obtained after permutation, identifying the same critical region as the sib-TDT, but was more extensive by spanning A3:119.1 – 129.4 Mb. Because the presentation has an autosomal recessive mode of inheritance, homozygosity mapping was also conducted and the same region was identified, further reducing the critical region to 2.1 Mb.

A variant dataset from WGS of domestic cats, the 99 Lives Cat Genome Sequencing Initiative, is available and has revealed the causative variants for several cat diseases and traits in the last several years [30,39-45]. WGS was performed on a trio, which included a carrier dam, an affected sire, and an affected offspring, and led to the identification of a candidate variant. Although *GDF7* c.221_227delGCCGCGC (p.Arg74Profs*17) was initially identified as a heterozygous allele in the affected offspring, visual inspection with IGV suggested the affected offspring was instead very likely homozygous for the variant, which was also validated with sanger sequencing. In addition, *GDF7* c.221_227delGCCGCGC was unique to the trio and was identified within the critical region. *GDF7* covers the region of chromosome A3:126,998,247 – 127,002,459 and within the identified critical region. The variant also segregated concordantly in the established pedigree and was not present in any cats unrelated to the pedigree (i.e., other 192 cats in the feline WGS dataset).

In humans, HPE is the most common malformation of the prosencephalon, and its prevalence is approximate 1 in 10,000 births [46]. A common feature of HPE includes the incomplete separation of the anterior part of the forebrain or telencephalon. The previous study indicated this feline heritable brain malformation syndrome resembled a mild form of HPE [20]. Many genes have been reported to cause HPE in humans (Reviewed in [47-49]). However, *GDF7*, also known as *bone morphology protein 12* (*BMP12)*, has not been reported to be associated with HPE in humans. Initially, GDF7 activity was shown to be required for the specification of neuronal identity in the spinal cord [50]. *GDF7* mRNA is expressed within the roof plate when commissural axons initiate to grow ventrally-directed. Furthermore, *GDF7*-null mutant mice show hydrocephalus, and they show considerable variation in the location of the dilated ventricle [50]. This evidence supports these findings that the frameshift mutation in *GDF7* causing the truncated protein is highly likely to be associated with this heritable brain malformation syndrome in cats.

The variable severity of the malformation in the cat pedigree was reported previously [20]. In humans, heterogeneity in familial HPE is also identified even if different individuals are carrying the same mutation [51-53]. The influence of environmental or teratogenic factors or modifier genes have been suggested for the spectrum (Reviewed in [46,47,49]). Assuming no exposure to teratogen and relatively homogeneous living environment, the presence of modifier genes is suspected for the variable severity of the dilated ventricles and supratentorial cysts in cats presented here.

In conclusion, the combination of GWAS, homozygosity mapping, and WGS identified a 7-bp deletion in *GDF7* (c.221_227delGCCGCGC) that is the most likely variant causing feline forebrain commissural malformation concurrent with ventriculomegaly and interhemispheric cysts in this domestic cat lineage. Furthermore, this study highlights the importance of *GDF7* in the neurodevelopmental course in cats and brings new insight into neurodevelopmental biology. Cat breeders can now perform a genetic test to eradicate the *GDF7* mutation from the breeding population.

## Supplementary Materials

The following are available online at [URL should be provided here before the publication.]. Table S1: Regions of homozygosity in cats with the inherited brain malformation syndrome. Figure S1: Pedigree of cats segregating for an autosomal recessive forebrain malformation. Figure S2: Multi-dimensional scaling plot and quantile-quantile plot of cases and controls for genome-association analyses. Figure S3: Haplotype analysis of cases and controls for a heritable forebrain malformation in cats. Figure S4: Protein sequence alignment of GDF7 in cats (*Felis catus*) and other species. Figure S5. Variant validation by Sanger sequencing and fragment analysis. File S1: Ped file for PLINK of cats genotyped using Illumina Infinium Feline 63K iSelect DNA Array. File S2: SNPs (n = 92) forming common haplotype for cats in the association studies.

## Acknowledgements

This study was supported by in part by NIH Office of Research Infrastructure Programs (OD R24OD01092), Winn Feline Foundation (MT-13-010), the Cat Health Network (D14FE-552) and the MU Gilbreath McLorn Endowment for Comparative Medicine (LAL). The authors thank the JSPS Overseas Challenge Program for Young Researchers (2017–2018) for sponsoring the visiting scholarship (YY) and the financial support from Mars, Inc (RMB). We also thank Barbara Gandolfi, PhD and Thomas R. Juba, MS, for technical assistance.

## Author Contributions

Conceptualization, L.A.L.; Methodology, L.A.L.; Software, R.M.B., Y.Y.; Validation, Y.Y.; Formal Analysis, E.K.C., Y.Y.; Investigation, E.K.C., Y.Y., R.M.B.; Resources, L.A.L., E.K.C.; Data Curation, L.A.L., R.M.B.; Writing – Original Draft Preparation, Y.Y., E.K.C; Writing – Review & Editing, Y.Y., R.M.B., E.K.C., L.A.L.; Visualization, Y.Y., R.M.B., E.K.C.; Supervision, L.A.L.; Project Administration, L.A.L.; Funding Acquisition, L.A.L.

## Conflict of Interest

Authors disclose no conflict of interest. The funding sponsors had no role in the design, execution, interpretation, or writing of the study. The authors may receive supportive funds from a genetic testing laboratory that would offer this variant as a commercialized test in the future.

---

**Figure S1.**
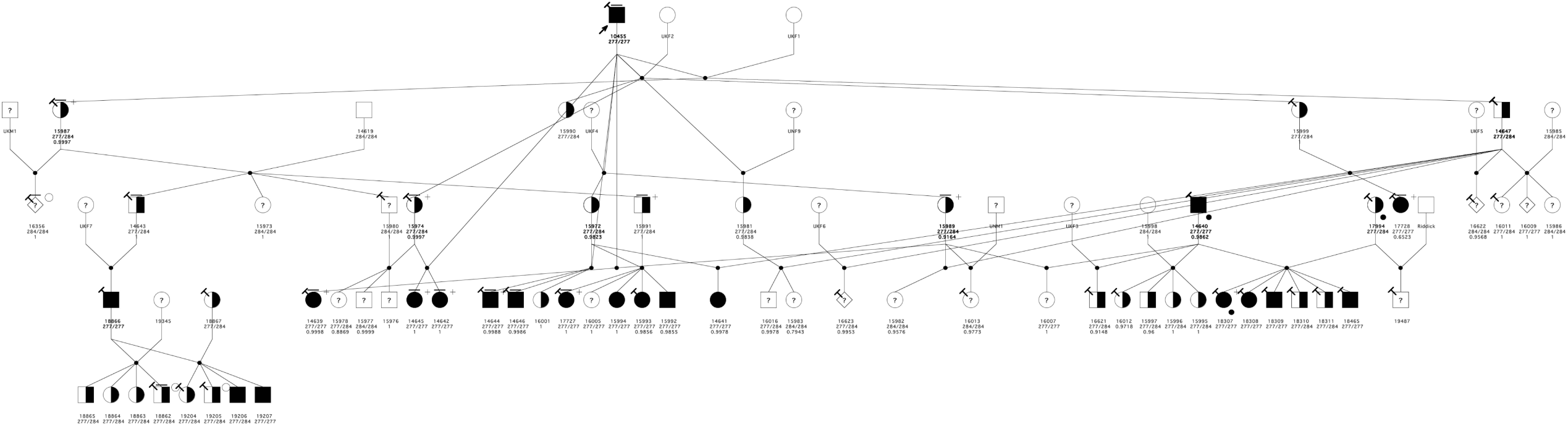
Pedigree of cats segregating for an autosomal recessive forebrain malformation. Relationships of 79 cats (27 nuclear families) provided by the breeder and confirmed with genetic testing of short tandem repeats when possible. Arrow indicates the proband. Circles indicate females, squares indicate males, and diamonds indicate unknown sex. Filled symbols represent cats with small rounded ears, which were suspected to have forebrain commissural malformation concurrent with ventriculomegaly and interhemispheric cysts. Half-filled represent obligate carriers. Symbols with question marks represent cats with unknown phenotype. A symbol with no fill indicates the cat is known to be completely unrelated and not expected to be a carrier. The cats genotyped on the DNA array and used for genome-wide association studies and homozygosity mapping are indicated by a “T” on the upper left of the symbol (The nine cats removed by quality control are not indicated). A black filled circle at the left bottom of symbol are individuals that were whole genome sequenced. Cats with a bar above the symbol were confirmed by magnetic resonance imaging. Cats with an open circle to the upper right had histology performed at necropsy. The cats’ ID/name is indicated below the symbol. Size in basepairs of the genotypes for the 7 bp *GDF7* indel are indicated below each cat available.

**Figure S2:**
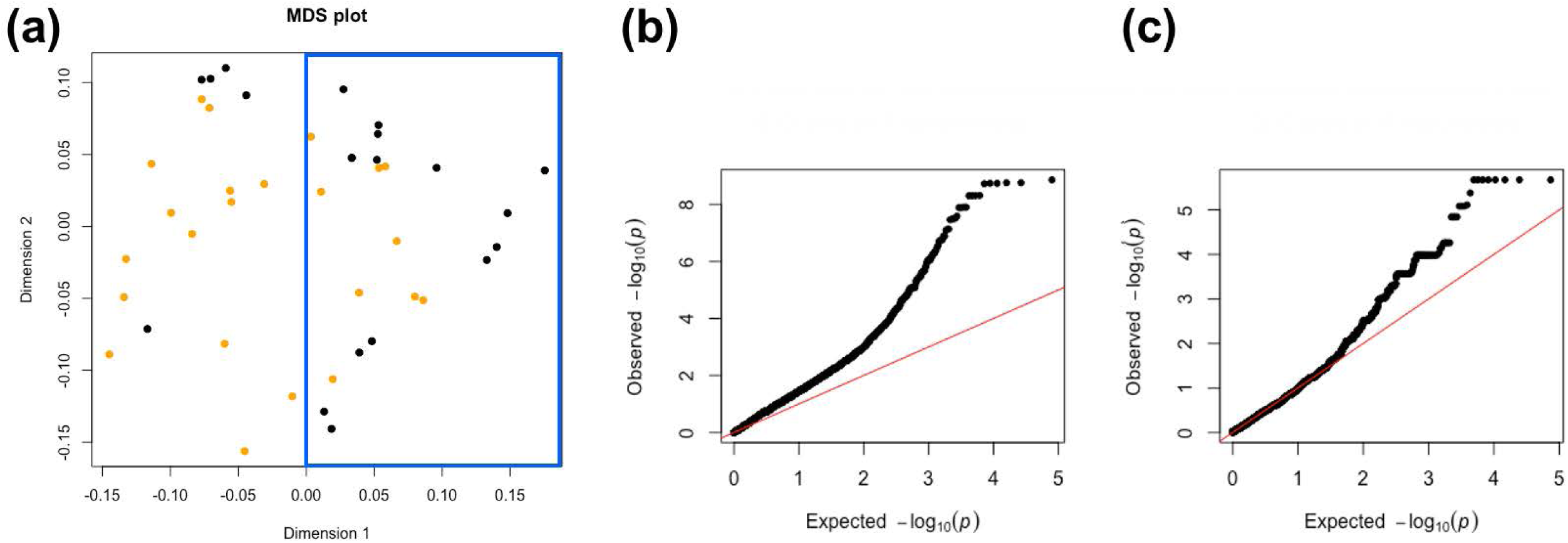
Multi-dimensional scaling plot and quantile-quantile plot of cases and controls for genome-association analyses. (a) Multi-dimensional scaling (MDS) plot of cats used for the initial case-control association analysis. The genomic inflation was 1.89. Therefore, cats clustered within the blue rectangular area were used for the second case-control association analysis. The genomic inflation factor was reduced to 1. (b, c) The quantile-quantile plots of cats used for the initial (b) and second (c) analyses demonstrate the observed versus expected–log(p) values.

**Figure S3.**
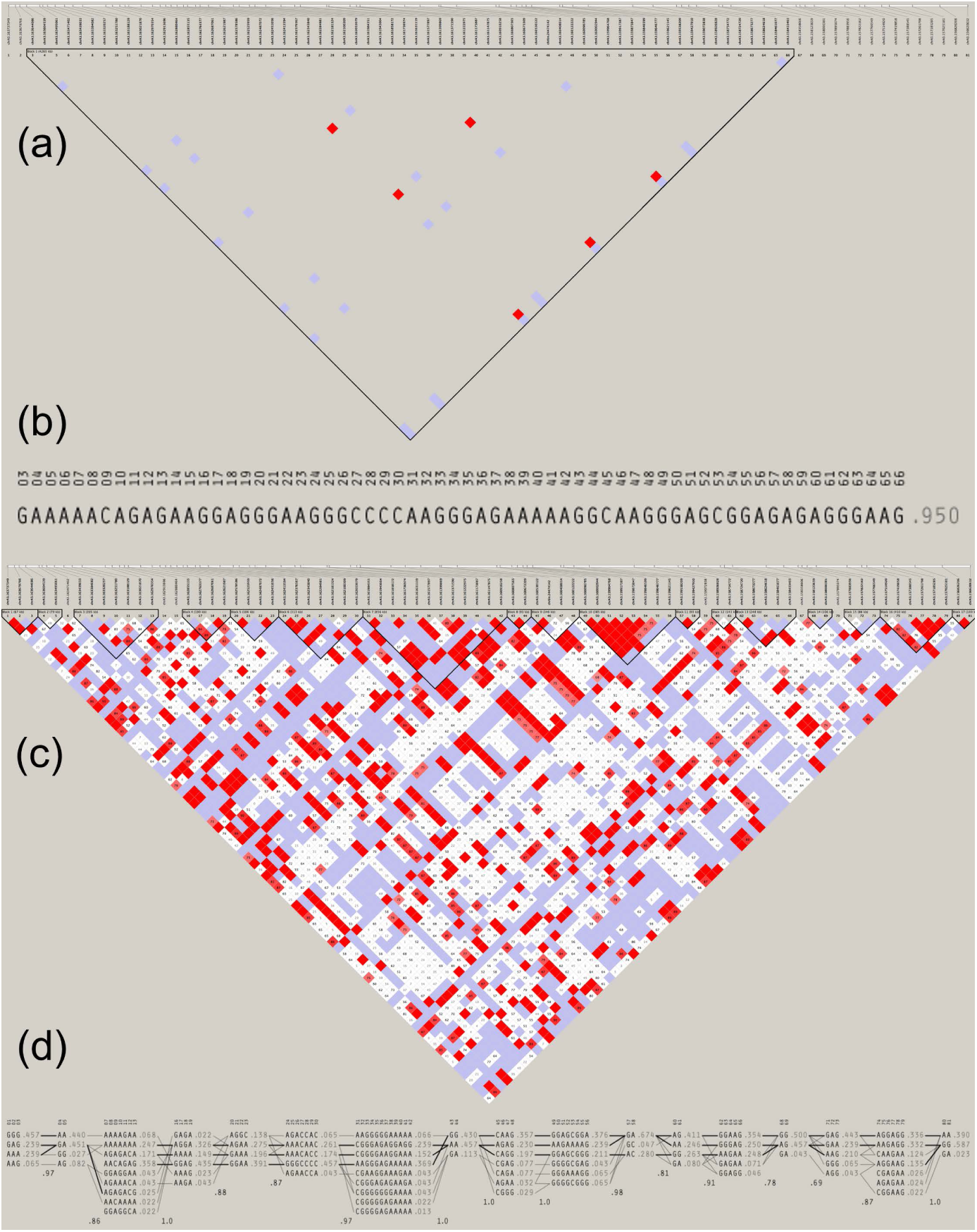
Haplotype analysis of cases and controls for a heritable forebrain malformation in cats. DNA array genotypic data for cats (20 cases and 23 controls) were analyzed for a common haplotype. Position 123,014,546 (chrA3.163737349) to position 128,837,125 (chrA3.156620632) on chromosome A3 encompassed 81 SNPs. (a) A large extended linkage disequilibrium block is identified by Haploview in cases, spanning 4,265 kb regions. (b) Sequential haplotype and its frequency within the cases. One major haplotype is shown and accounts for 95% in all cases. The remaining 5% haplotypes are not listed because each of those haplotypes has a frequency of <1% (c) Short and discontinuous linkage disequilibrium blocks are identified by Haploview in controls. (d) There are various haplotype sequences and frequencies within approximately 6 Mb regions in unaffected cats.

**Figure S4.**
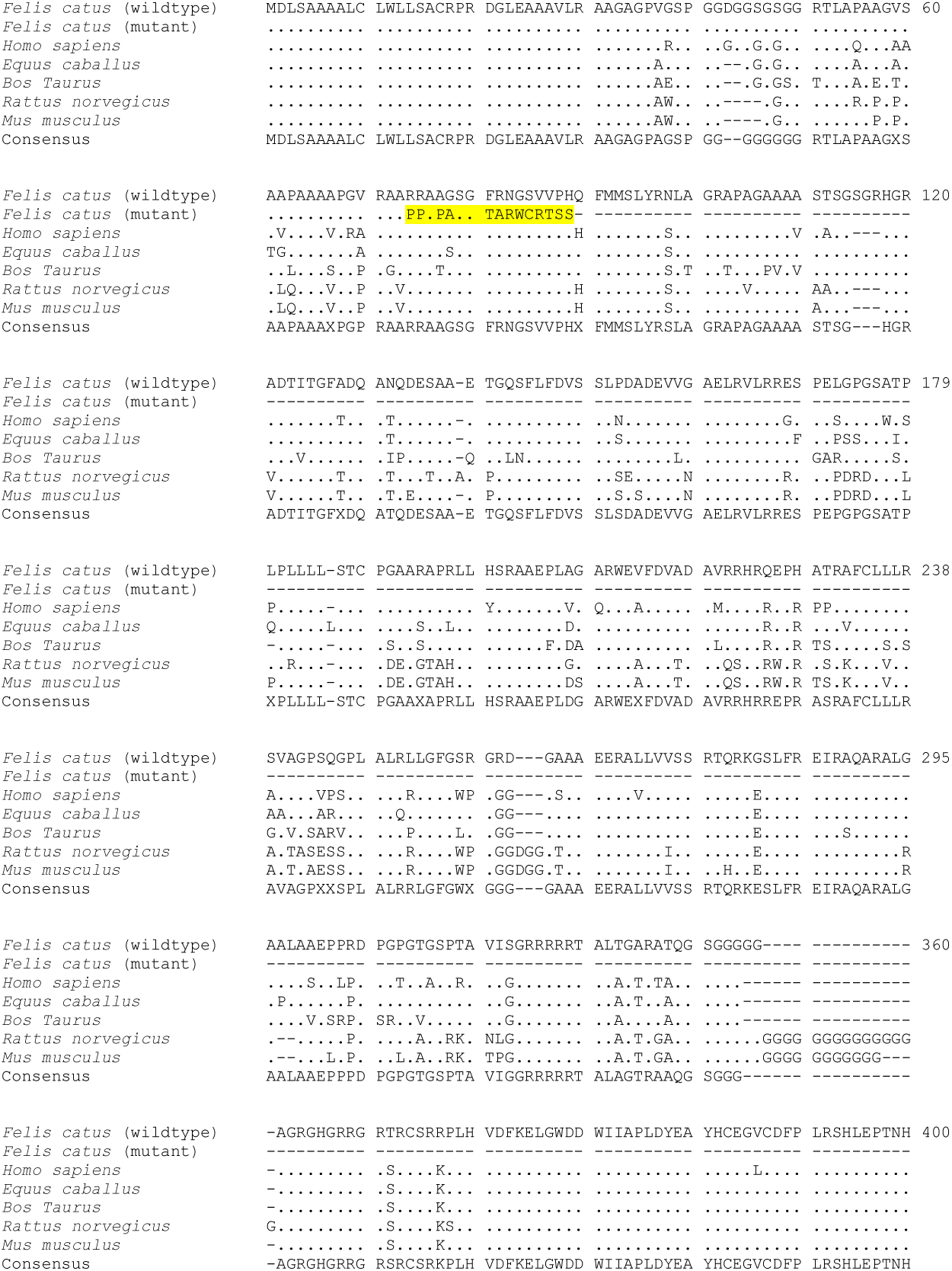
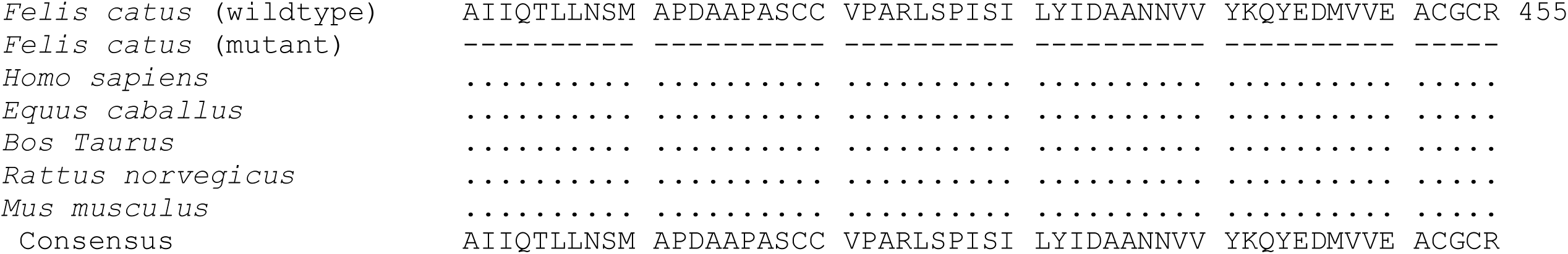
Protein sequence alignment of *GDF7* in cats (*Felis catus*) and other species. GDF7 protein sequences are aligned from wildtype cat (*Felis catus*), *GDF7* mutant cat, cow (*Bos Taurus*: NP_001193030.1 [ARS-UCD1.2]), horse (*Equus caballus*: XP_023475218.1 [EquCab3.0]), mouse (*Mus musculus*: NP_001299805.1 [GRCm38.p4]), and rat (*Rattus norvegicus*: XP_006239940.1 [Rnor_6.0]). Identical amino acids to those of *Felis catus* sequence are represented as a dot (.). Deleted amino acids are represented as a dash (–). A 7 bp deletion causes a frameshift and changes the amino acid sequence from 74th position (highlighted in yellow), starting with an arginine to a proline change, which results in the truncated protein with a stop codon 17 amino acids downstream.

**Figure S5.**
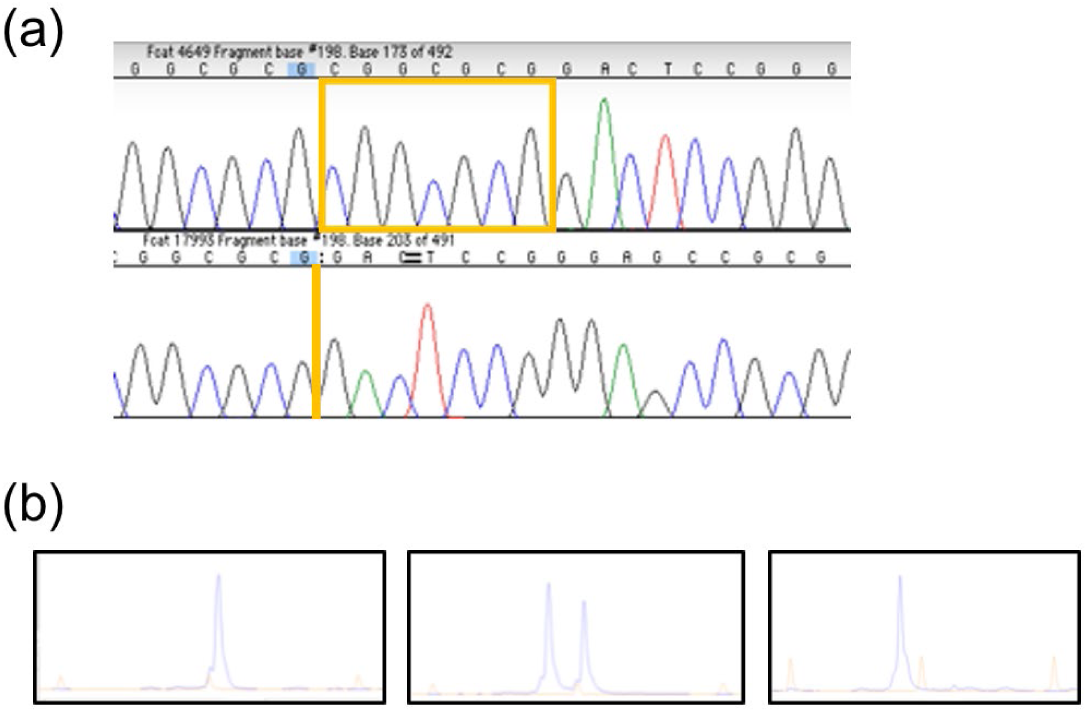
Variant validation by Sanger sequencing and fragment analysis. Sanger sequence of a wildtype and homozygous affected cat for the 7 bp *GDF7* variant (boxed region). (B) Fluorescence-based fragment analysis using an ABI 3730XL for the *GDF7* variant. Left – homozygous wildtype with 294 bp fragment, middle – heterozygous with 287 and 294 bp fragments, and right – affected with 287 bp fragment. LIZ standard (Applied Biosystems, Foster City, CA, USA).

**Table S1.**
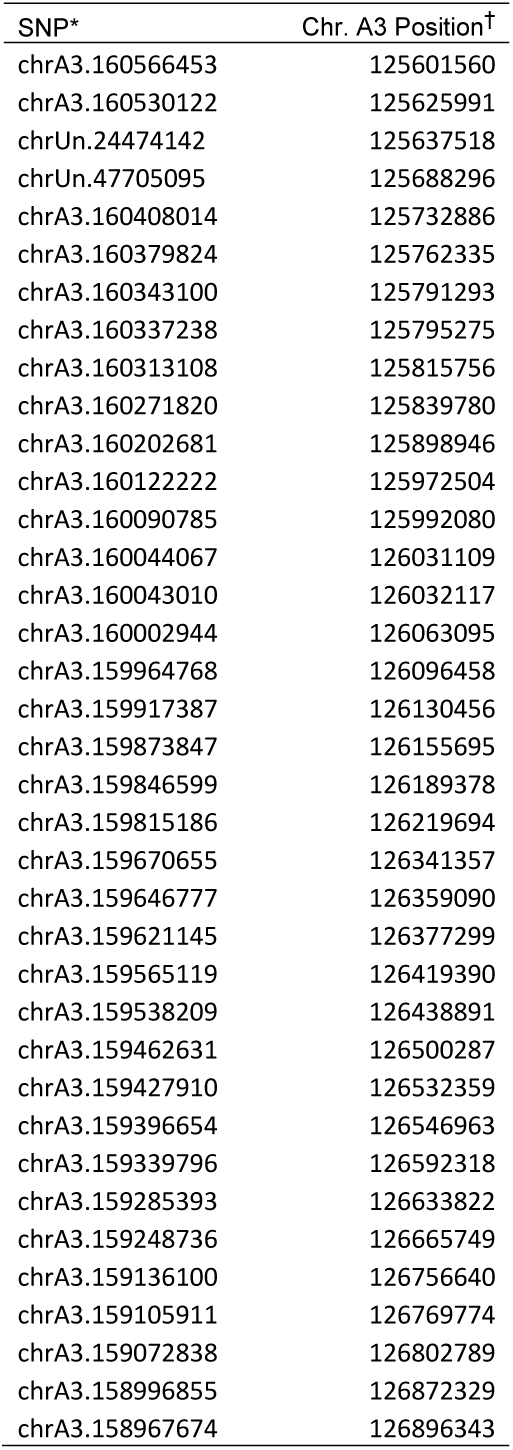
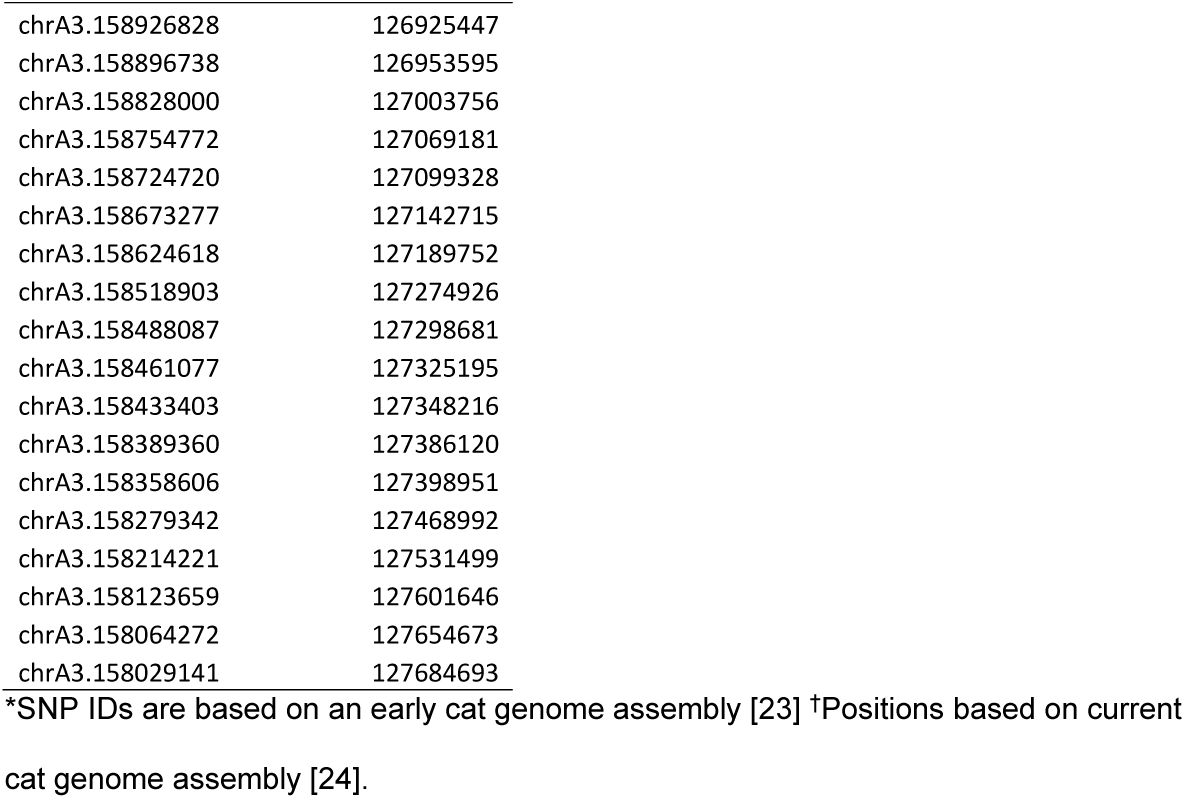
Regions of homozygosity that was unique to 18 cats with the inherited brain malformation syndrome, and was absent in all the unaffected cats.

| SNP* | Chr. A3 Position<sup>†</sup> |
| --- | --- |
| chrA3.160566453 | 125601560 |
| chrA3.160530122 | 125625991 |
| chrUn.24474142 | 125637518 |
| chrUn.47705095 | 125688296 |
| chrA3.160408014 | 125732886 |
| chrA3.160379824 | 125762335 |
| chrA3.160343100 | 125791293 |
| chrA3.160337238 | 125795275 |
| chrA3.160313108 | 125815756 |
| chrA3.160271820 | 125839780 |
| chrA3.160202681 | 125898946 |
| chrA3.160122222 | 125972504 |
| chrA3.160090785 | 125992080 |
| chrA3.160044067 | 126031109 |
| chrA3.160043010 | 126032117 |
| chrA3.160002944 | 126063095 |
| chrA3.159964768 | 126096458 |
| chrA3.159917387 | 126130456 |
| chrA3.159873847 | 126155695 |
| chrA3.159846599 | 126189378 |
| chrA3.159815186 | 126219694 |
| chrA3.159670655 | 126341357 |
| chrA3.159646777 | 126359090 |
| chrA3.159621145 | 126377299 |
| chrA3.159565119 | 126419390 |
| chrA3.159538209 | 126438891 |
| chrA3.159462631 | 126500287 |
| chrA3.159427910 | 126532359 |
| chrA3.159396654 | 126546963 |
| chrA3.159339796 | 126592318 |
| chrA3.159285393 | 126633822 |
| chrA3.159248736 | 126665749 |
| chrA3.159136100 | 126756640 |
| chrA3.159105911 | 126769774 |
| chrA3.159072838 | 126802789 |
| chrA3.158996855 | 126872329 |
| chrA3.158967674 | 126896343 |
| chrA3.158926828 | 126925447 |
| chrA3.158896738 | 126953595 |
| chrA3.158828000 | 127003756 |
| chrA3.158754772 | 127069181 |
| chrA3.158724720 | 127099328 |
| chrA3.158673277 | 127142715 |
| chrA3.158624618 | 127189752 |
| chrA3.158518903 | 127274926 |
| chrA3.158488087 | 127298681 |
| chrA3.158461077 | 127325195 |
| chrA3.158433403 | 127348216 |
| chrA3.158389360 | 127386120 |
| chrA3.158358606 | 127398951 |
| chrA3.158279342 | 127468992 |
| chrA3.158214221 | 127531499 |
| chrA3.158123659 | 127601646 |
| chrA3.158064272 | 127654673 |
| chrA3.158029141 | 127684693 |

\*SNP IDs are based on an early cat genome assembly [23] †Positions based on current cat genome assembly [24].

## Notes

### Competing Interest Statement

The authors have declared no competing interest.

